# *In-silico* investigation of umami peptides with receptor T1R1/T1R3 for the discovering potential targets: A combined modeling approach

**DOI:** 10.1101/2021.10.10.463792

**Authors:** Wenli Wang, Zhiyong Cui, Menghua Ning, Tianxing Zhou, Yuan Liu

## Abstract

Umami, providing amino acids/peptides for animal growth, represents one of the major attractive taste modalities. The biochemical and umami properties of peptide are both important for scientific research and food industry. In this study, we did the sequence analysis of 205 umami peptides with 2-18 amino acids, sought the active sites of umami peptides by quantum chemical simulations and investigated their recognition residues with receptor T1R1/T1R3 by molecular docking. The results showed the peptides with 2-3 amino acids accounting for 44% of the total umami peptides. Residues D and E are the key active sites no matter where they in peptides (N-terminal, C-terminal or middle), when umami peptides contain D/E residues. N69, D147, R151, A170, S172, S276 and R277 residues in T1R1 receptor were deem to the key residues binding umami peptides. Finally, a powerful decision rule for umami peptides was proposed to predict potential umami peptides, which was convenient, time saving and efficiently.

## 1. Introduction

Umami as an alimentary taste not only enhances food palatability, but also serves as an indicator of nutrients for human body[1]. Umami has been listed in the basic taste after the discovery of umami receptors[2]. The current umami ligands contain, but are certainly not limited to free L-amino acids, purine nucleotides, peptides, organic acids, amides and their derivatives[3]. Thereinto, a few umami peptide molecules from hydrolysates of fish protein or other foods have become a new interest with the development of umami agent based on the especial umami characteristics of peptides[4]. In recent years, more peptides have been reported showing umami (enhancing) taste[5–7]. Due to the multivariate taste characterization (umami, sweet, sour, salty and kokumi) of peptides (Supplementary table 1), the extent of the umami mechanism of peptides is not fully understood, therefore, umami peptides are questioned for taste characteristics. The taste mechanism and characteristics of peptides have only begun to be appreciated recently. Umami peptides can be predicted and analyzed by using a scoring card method with propensity scores of dipeptides[8]. The structure features of umami hexapeptides were analyzed by three-dimensional quantitative structure-activity relationship (3D-QSAR) modeling[9].

Umami is induced by the ligands binding to T1R1-T1R3 heterodimers of taste receptor type 1 (T1R)[2] that can recognize a wide array of umami taste emerged by amino acids, peptides, nucleotides through its extracellular ligand-binding domain (LBD)[10]. The amino acid sequences and spatial structures of umami substances are the key to umami recognition based on the studies of umami receptors[10, 11]. To obtain a more comprehensive understanding of umami recognition mechanism, homology modeling, molecular docking and molecular dynamics simulation were applied to construct a computational modeling of T1R1-T1R3 heterodimers due to no crystal structure[12, 13]. Although the binding mode of glutamate with T1R1 has been intensively studied[14, 15], the extent of umami peptides recognition mechanism has only recently begun to be appreciated. Examples including three hexapeptides from Atlantic cod[16], five umami peptides from myosin[17] and a GK-15 peptide from tempeh[18], which can dock with the principal amino acid residues of T1R1/T1R3, showed umami taste by sensory evaluation.

Umami peptides are of comparable importance in all umami ingredients. Knowledge of the diversity of known umami peptides is essential for understanding their umami rule. To elucidate the activation mechanism of umami peptides on receptors, this study focuses on the know umami peptides (Supplementary table 1) in March 2021 data (Web of Science, database). We analyzed the amino acid information of umami peptides, investigated the chemical active sites of umami peptides and the putative binding modes with the hT1R1 receptor. According to the sequence and active site information of peptides, we proposed a novel decision rule for umami peptides to valuate those potential umami peptides.

## 2. Materials and Methods

### 2.1 Chemical activity analysis of peptides based on quantum chemical simulations

The article analyzed the taste mechanism of peptides at the molecular level. Molecular modeling started with the construction of the identified umami peptides using GaussView 5.0.9, and all computations were carried out using the Gaussian 09 program. The identified umami peptides were optimized using density functional theory method (DFT) in theory level B3LYP/6-311G (d, p), and the frequencies were also calculated with the same method to ensure the minimum energy structure [19]. Subsequently, the Molecular electrostatic potential (MEP)’s generated from the atomic charge. The frontier orbitals (HOMO and LUMO) were calculated with the aid of Molekel program. Under this strategy, it was possible to obtain the best potential quantum molecular series points defined around the molecule. These illustrated the chemical information of the molecule or the potential active site that easily reacts with special enzyme/receptors.

### 2.2 Molecular Docking

In order to further explain the umami characteristics of the above peptides, hT1R1-T1R3 VFD model based on the fish T1R2a-T1R3 structure (PDB id:5×2M) was used in this study[10]. The optima structures of umami peptides were constructed using Gauss-View 5 and Amber 14 software. Due to the auxiliary role of hT1R3 for ligand binding[15, 20], the molecular docking about the above peptides to hT1R1-VFD was performed using AutoDock vina-12 software package with the default scoring function. The PDBQT parameters of receptor and ligands were calculated by Autodock Tools-1.5.6 package with the docking box-size 30 and the box center as the ligand center.

## 3. Result and Discussion

### 3.1 The amino acid composition profiles of known umami peptides

The previous studies showed that acidic amino acid (aspartic acid (D) and glutamic acid (E)) were generally contained in the sequence of umami peptides[4]. Alanine (A), serine (S), glycine (G) as the sweet-taste amino acids can synergize with D/E and elicit to umami taste[3, 21]. The umami characteristics of peptides are related to their primary sequences. Moreover, Phasit et. al designed SCM-derived propensity scores to predict and analyze the umami intensities of peptides based on the dipeptide in sequence information[8]. Knowledge of the primary sequence information of each umami peptide was essential for understanding their umami characteristics. To fully investigate the amino acid composition profiles of umami peptides, we summarized a total of 205 umami peptides with 2-18 amino acids (Fig. 1A). We found that 91 predominant umami peptides containing 2 or 3 amino acids accounted for 44% of total umami peptides. Besides, hexapeptide and octapeptides also accounted for 11% and 9% of total umami peptides, respectively. The umami peptides with more than 10 amino acids have not been intensively reported, and the extent of which these peptides have only recently begun to be appreciated[7, 18, 22].

**Fig. 1.**
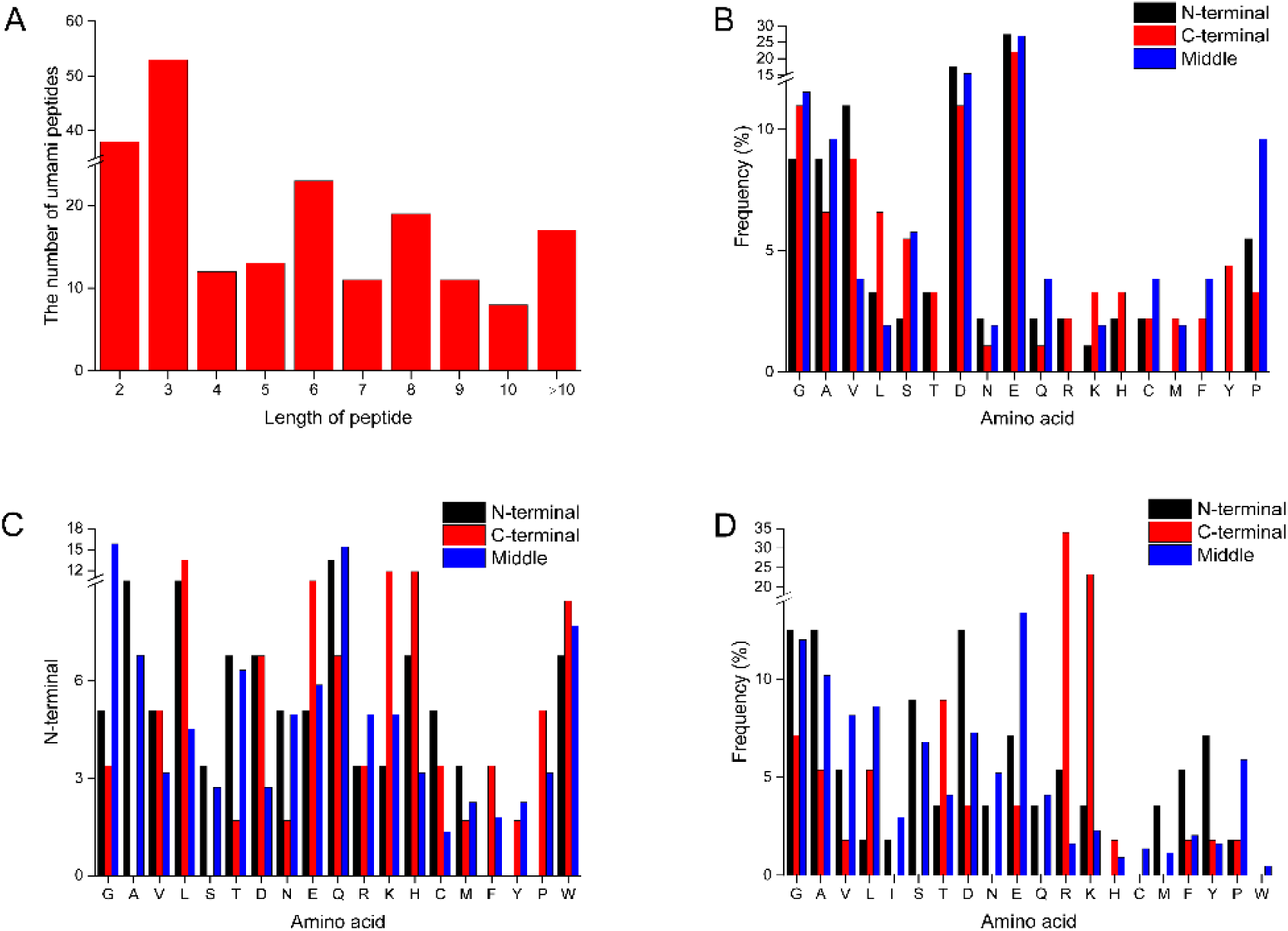
Amino acid composition profiles of umami peptides analysed in this work. A: The peptide-length distribution of umami peptides; B: The peptide with 2-3 amino acids; C: The peptide with 4-7 amino acids; D: The peptide with >7 amino acids

Considering the potential role of peptide length in umami characteristics, 205 umami peptides were divided into three types according to the peptide length and discussed based on the frequency of residues in peptides. For the umami peptides containing 2 or 3 amino acids, residue D and E are the predominant residues no matter at any position (N-terminal, C-terminal or middle) (Fig. 1B). Results of the proportion of D & E in these peptides were in consistent with the definition of D/E as umami related amino acids[4]. In addition, we also found other residues that were significantly exceeded, such as residues G, A and V at middle position. About 30% of the known umami peptides have 4-7 amino acids. Among these peptides, compared with D and E the high frequency residues were biased toward residues L, Q, and K (Fig. 1C), which was a striking departure from the discovery of umami di-/tripeptides. Residues L, D and K all showed higher frequency in the C-terminal of umami peptide. However, residues A, S, T and N are not found as C-terminal residues, and residues F, Y and P as N-terminal residues are not found in these umami peptides. Last but not least, we also found that residue W showed high frequency in the umami peptides with 4-7 amino acids.

In contrast to the peptides with 4-7 amino acids, the amino acid profiles of the peptides with >10 residues were significantly different from the distribution in small peptides. Though their amino acid distribution did not show an obvious rule, residues R and K as the high frequency residues accounted for 20-35% of total C-terminal residues (Fig. 1D). Previous studies have also provided evidence that the presence of residue R as C-terminal amino acid residue affected on the taste intensity of umami peptides[7]. Residue G at any position all showed high frequency, which may be related to their own features as the sweet-taste amino acids. In addition, residue A, S and D all have a certain frequency (over 10%) in the N terminal in long peptides. In summary, based on the amino acid investigation of umami peptides, we could understand and explain the amino acid distribution in umami peptides.

### 3.2 The key active site in umami peptides

It is well known that the frontier molecular orbitals (FMOs) play a key role in many aspects of a compound, for example electric properties[23]. In the current study, the potential active sites in 205 peptide molecules were identified according to the calculation of HOMO-LUMO. Among these peptides, the calculation of 17 umami peptides was failed due to the particularity of their spatial structure, and we obtained the active site results from 188 peptides. For the umami peptides with 2-3 amino acid residues, one of D, E, R or K residue can be observed as the key active sites when umami peptide contains residue D or E. In support of this idea, D and E as the key umami amino acids have been known in most taste studies[3, 24]. With the absence of residue D or E in peptides, N, Q, G, A, S, T and H would be easier to become the active sites when residue L, F or I were absent in peptides (Table S1). However, a closer examination of our data provides interesting insights regarding the C-terminal of peptide. It does suggest that C-terminal residues contribute to more active sites than N-terminal residues. Surprisingly, R or K as the C-terminal residues always are the active sites of peptides no matter whether there are residue D & E or not.

The distributions of active sites in the umami peptides with 4-7 amino acid residues were mostly in agreement with those of di-/tripeptides. Residue D or E as the key active sites were still found in those umami peptides which contain D or E residue. The rule that C-terminal residues contribute to more active sites is also applicable to the umami peptides with 4-7 amino acid residues. If D or E residue presents, residue R or K would be the active sites together with D or E residue. The active sites of peptides are also biased toward R or K due to the absent of C-terminal residues of D or E residues. Compared with other amino acid residues, in the presence of D or E residue, Q, N, A, H and S showed a certain frequency at the active sites of peptides. In these umami peptides, Q, N, A, H and S still showed a certain activity in the absence of D or E residue when they also did not contain the hydrophobicity residue L, F, Y or I. It may be the reason why these amino acids always show bitter taste, because they mostly exist in bitter peptides[25, 26].

Although the active profiles of peptides with more than 7 amino acids residues are diversity owing to the complexity and inhomogeneity of amino acid composition, D or E was still the predominant active residue. Most R or K as the C-terminal residues were calculated as the active sites in umami peptide but those peptides with more than 10 amino acids were not included. The importance of D or E in umami peptides indicates their contribution in umami perception. In support of this idea, D and E have been previously reported as two typical critical umami amino acids^[4]^. Based on the above-mentioned distribution trait of active site in umami peptide, we obtained a propensity umami score of umami peptide (Fig. 2). The most contributing active sites (D or E) were set to 0.5 score, followed by active sites (R or K) as the score of 0.3, and active sites N, Q, G, A, S, T and H were set to score 0.2, separately. The umami scores of peptides were calculated by combining Score A with Score B. These score data were clustered into 4 groups by applying K-means clustering algorithm. In the umami peptides with 2-3 amino acids, about 70% peptides were classified in the cluster with score over 0.5, which indicated that they contained a key active site (D or E) or two active sites likes (R, K, N, Q, G, A, S, T) (Fig. 2A). The peptide with score >0.5 could be deemed as the higher potentiality umami peptide. We also analyzed the peptides with 4-7 amino acids as well as those containing more amino acids, and they all contained at least 56% of peptides in the cluster of with score >0.5 (Fig. 2B&C). Although this approach cannot identify exact contacts between amino acids and umami receptor, it as a unique and effective method does propose probable residues that contribute to umami perception. These data will serve as an important resource for further research on the rule of umami in peptides.

**Fig. 2.**
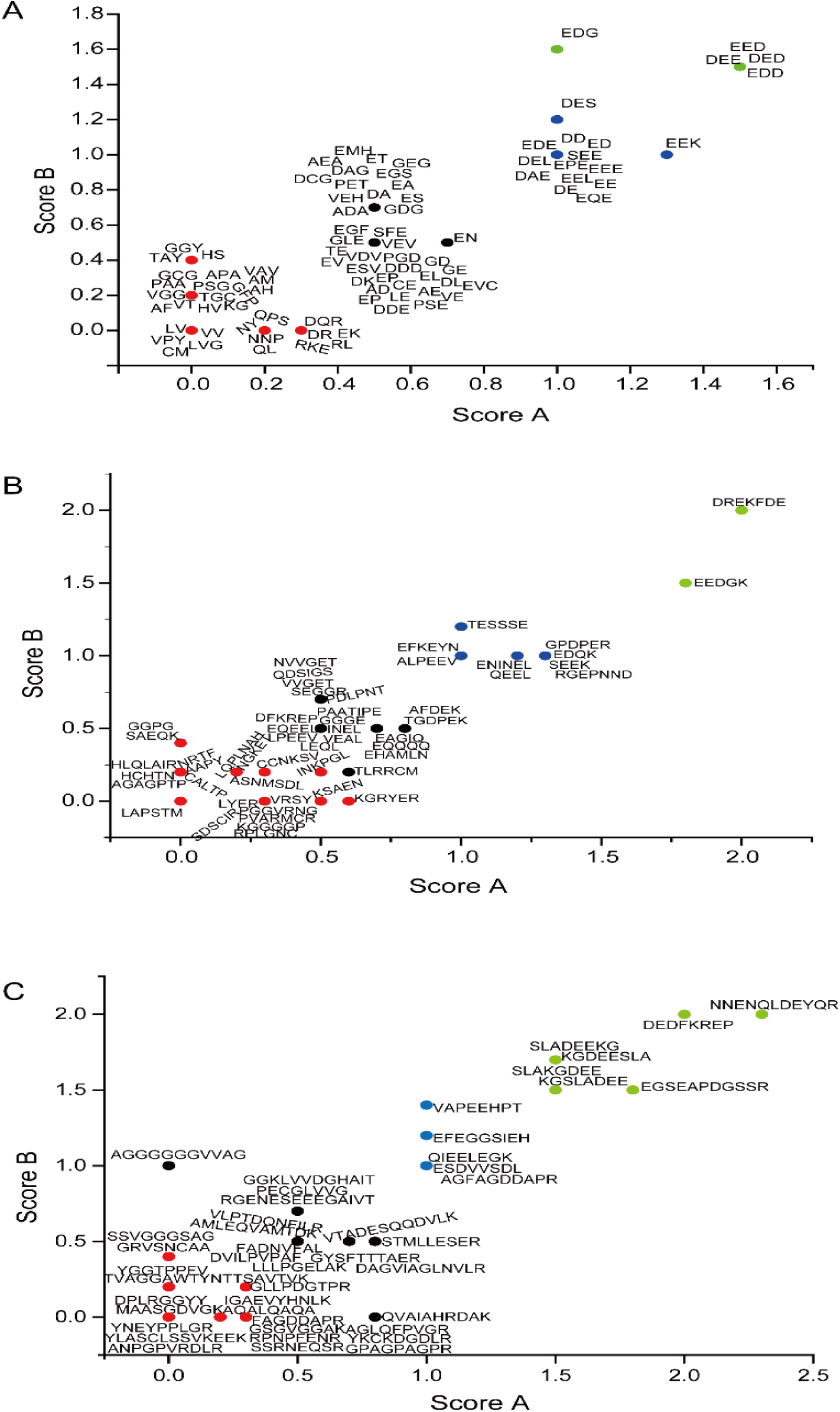
The propensity umami scores of umami peptides based on their active sites. Residue D and E were set separately a score of 0.5; residue R and K were set separately a score of 0.3; and residue N, Q, G, A, S, T and H were set separately a score of 0.2. Each peptide was calculated by the Score A combining with Score B. Thereinto, Score means the sum of 0.5*D+0.5*E+0.3*R+0.3*K+0.2*N+0.2*Q, and Score B means the sum of 0.5*D+0.5*E+0.2*G+0.2*A+0.2*S+0.2*T+0.2*H. A: Umami peptides with 2-3 amino acids; B: Umami peptides with 4-7 amino acids; C: Umami peptides with >7 amino acids

### 3.3 The rule of umami peptides-bind sites of T1R1

So far, eight candidate umami taste receptors have been reported^2^. Thereinto, heteromeric T1R1/T1R3 in response to a wide range of structurally diverse umami compounds, can be extensively regulated and activated by L-glutamic acid, disodium inosinate, disodium guanylate and disodium succinate based on the results of T1R1 receptor sensors[27]. The understanding of ligand binding to T1R1/T1R3 at molecular level currently relies on computational methods including mathematical modelling and molecular docking due to the lack of experimental structural information on T1R1/T1R3 [10, 28]. These techniques were also applied for finding potential sweet molecules, expediting drug discovery, identification and characterization of bitter peptides, and investigating cognation of bitter compounds with bitter taste receptors[26, 29–31]. To obtain a more comprehensive understanding of the molecular recognition mechanisms of known umami peptides, the binding modes of 205 umami peptides to T1R1/T1R3 receptor were investigated. The results of umami peptides mentioned above provided extensive data. To exactly understand their binding modes, we analyzed these data separately according to the length of peptide.

The shared binding sites including N69, D147, R277 and Q278 in T1R1 indicated the highly conserved residues of di-/tripeptides (Fig. 3A). Moreover, these key sites in T1R1 mainly bound to the active sites of peptides, such as residue D, E, G, S, A and V. Based on the frequency of T1R1 binding to the active sites of ligands, residues S48, H71, A170, S172 and A302 in T1R1 were also the shared binding sites, except for the four residues mentioned above (Fig. 4A). Therefore, the active sites of ligands are of comparable importance in understanding the binding modes of umami peptides with T1R1. Furthermore, we conducted analysis to attempt to identify the preferred binding site for the umami peptides with 4-7 amino acid residues by binding the shared binding sites in T1R1 and their frequency of binding to the ligand active sites. Notably, the set of peptides was biased toward residues N69, S147, N150, R151, A170, S276 and R277 in T1R1 (Fig. 3B & Fig. 4B). We found residues that were highly similar to those obtained in di-/tripeptides analysis including N69, D147, A170 and R277. For the set of peptides with >7 amino acids residues, these peptides involved more binding residues of T1R1 due to the complexity and multiformity of amino acid compositions. They had a highly significant selection to L51, N69, D147, R151, A170, S172, R277, A302 and S384, which does suggest that these residues contribute to substrate binding (Fig. 3C). We compared the frequency of active sites binding to each position in T1R1, especially for these high selection residues (Fig. 4C). We found that R151 in T1R1 was the predominant binding residue. Based on these key binding residues, the potential binding regions of the three sets of peptides to T1R1 were shown in Figure 5.

**Fig. 3.**
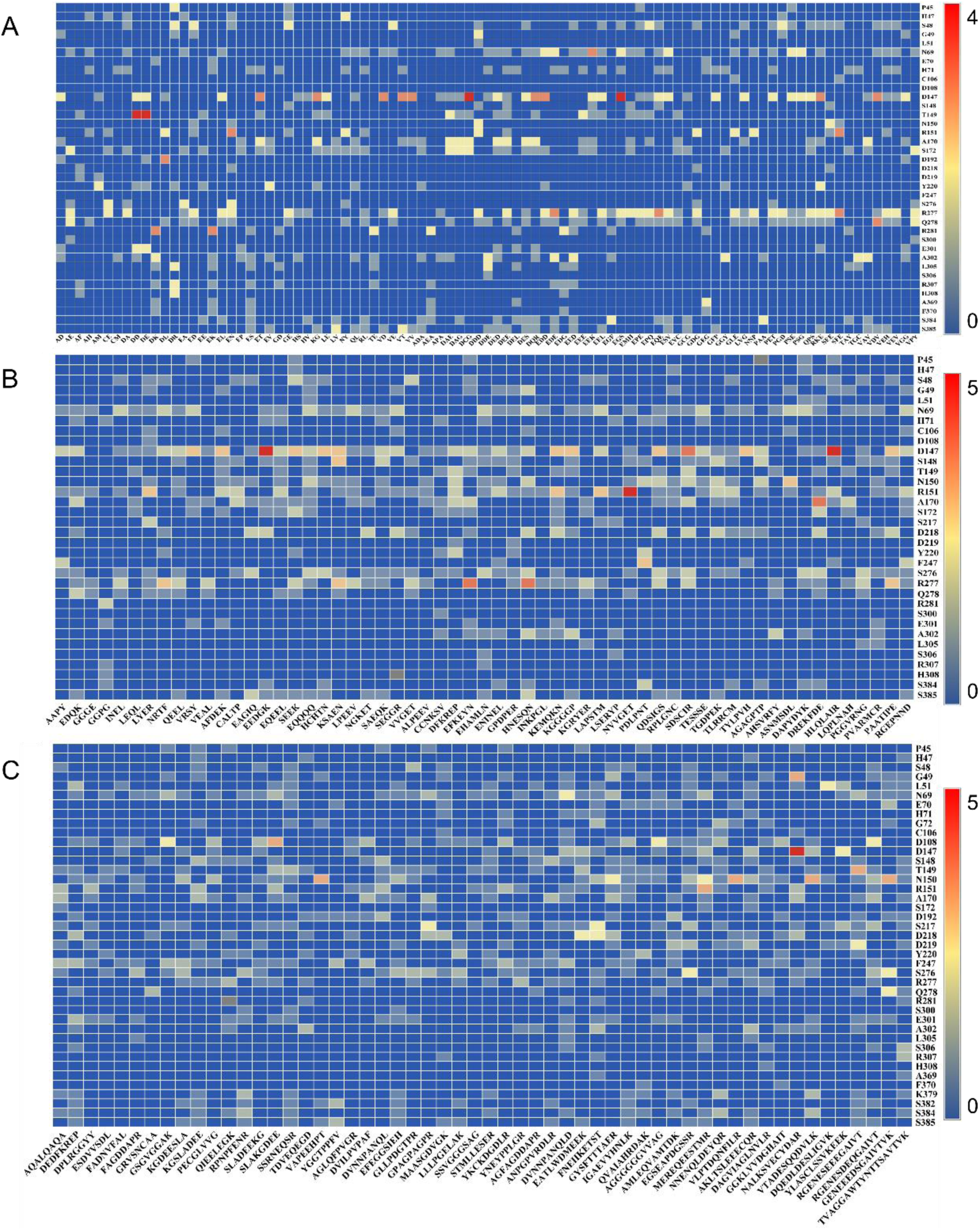
Docking sites of umami peptides into T1R1 binding pocket and their hydrogen bond values. A: Umami peptides with 2-3 amino acids; B: Umami peptides with 4-7 amino acids; C: Umami peptides with >7 amino acids

**Fig. 4.**
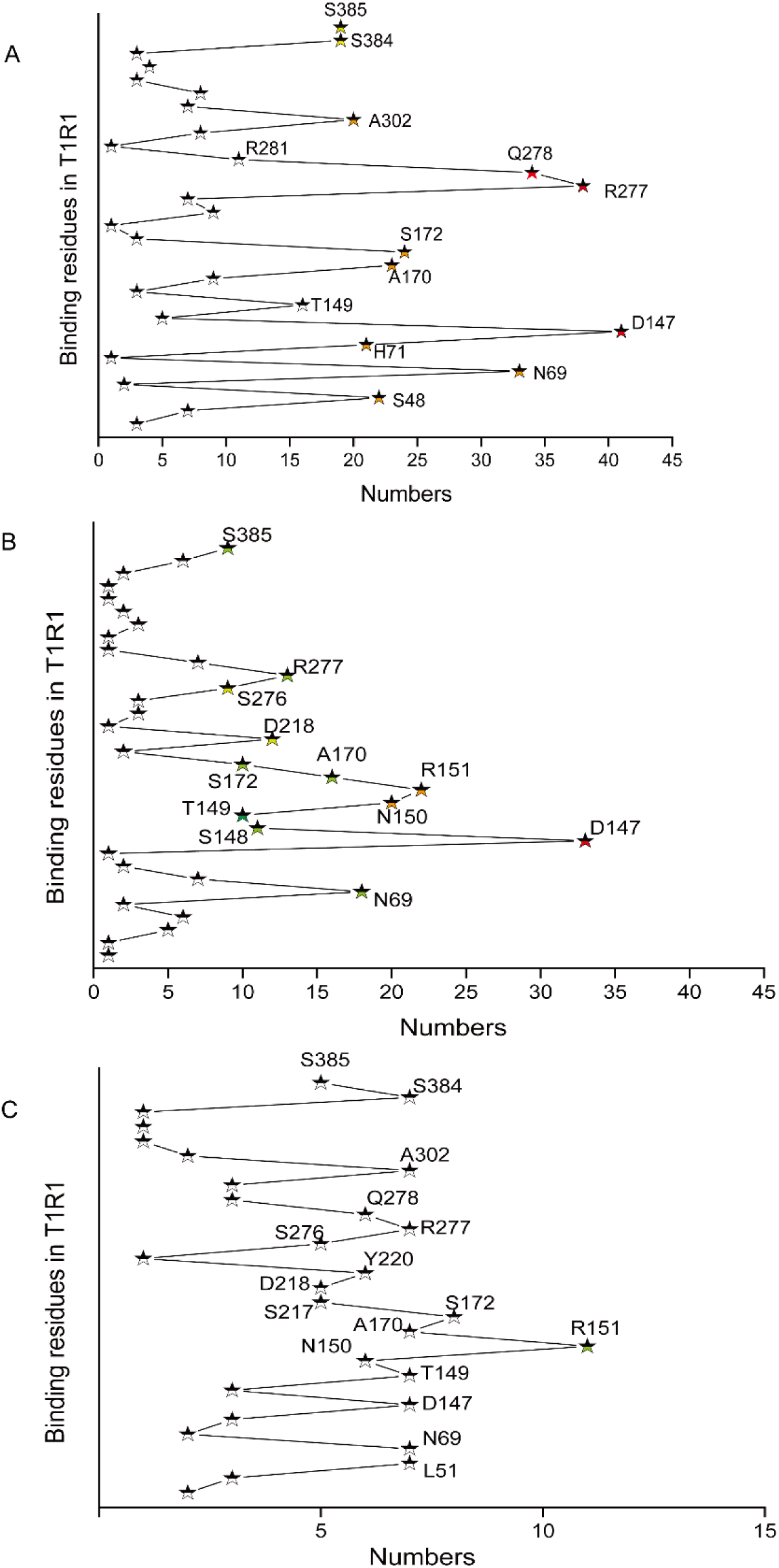
The number of active sites in umami peptides binding into T1R1 residues. A: Umami peptides with 2-3 amino acids; B: Umami peptides with 4-7 amino acids; C: Umami peptides with >7 amino acids

**Fig. 5.**
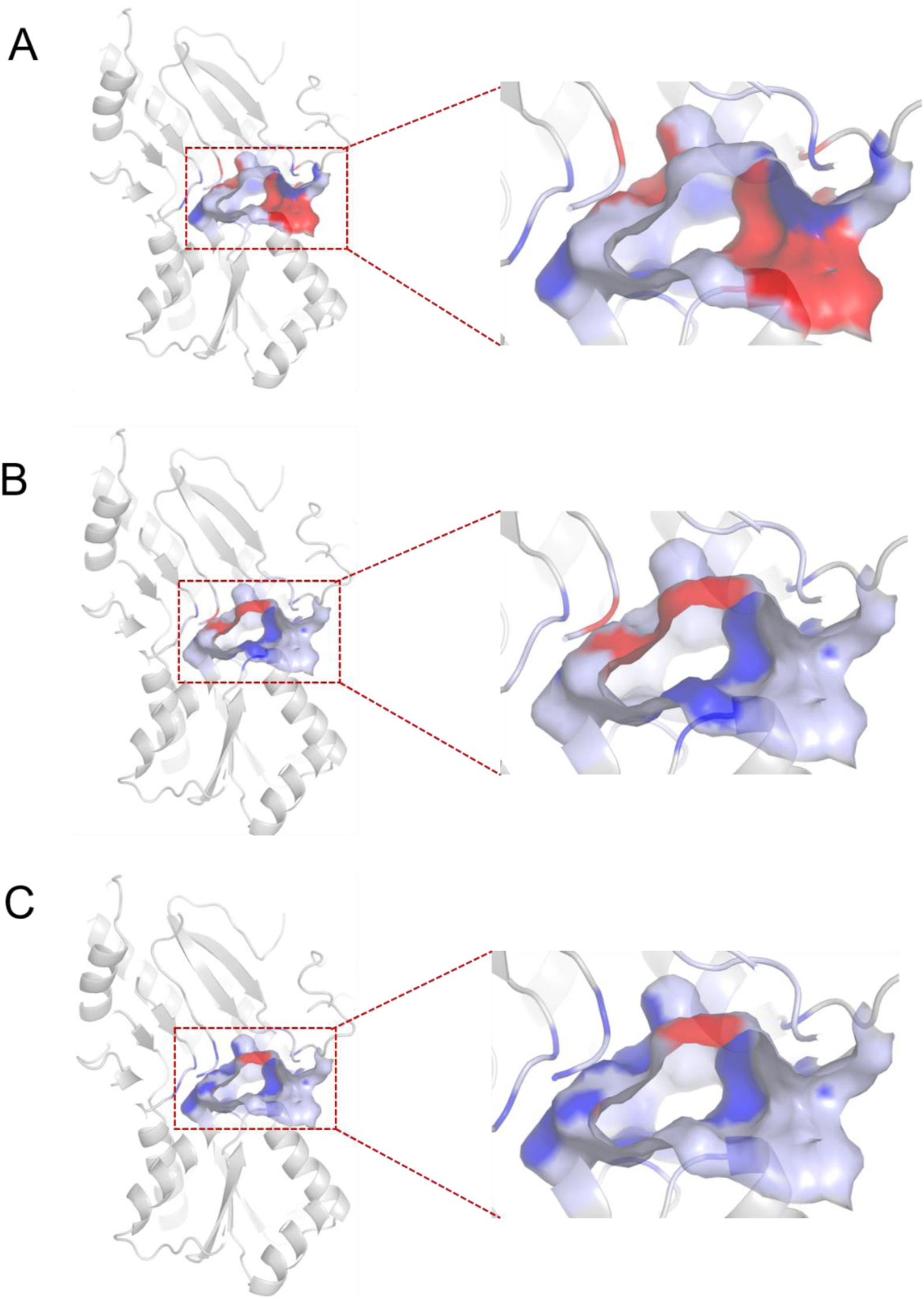
Residues defining the key binding site are colored in color, red and blue based on the docking results of umami peptides and the number of active sites in umami peptides binding into T1R1 residues

Closer examination of these data revealed an interesting similarity regarding the binding rule of umami peptide to T1R1. The highly conserved binding sites N69, D147 and R277 were found in the three sets of peptides. With the length of peptide increasing, more potential common binding sites were analyzed, such as N150, R151 S276 and A170. To support this idea, comparing our results to the studies of natural umami ligands showed interesting similarities. The effects of five natural umami ligands on the structural dynamics of T1R1-T1R3 were explained using molecular dynamics simulations[10, 28]. They identified that the key residues like N69, D147, R151, A170, S172, S276 and R277, were also consistent with our results ^[10]^, which suggested that they may be the common binding site for all umami ligands. It is particularly noteworthy that, the key finding from the above data is that the active sites of peptides including D, E, G, S, A and V is preferred to binding to receptor T1R1. This further proved that the active site in ligands is an important feature to explain the mechanism of umami taste.

### 3.4 Putative binding modes of umami peptides

Knowledge of umami peptides’ characterization is essential for understanding their taste mechanisms. However, no exact method has been developed to predict and characterize umami peptides based on active sites of umami peptides. In the current study, we propose a novel decision rule by combining the information of amino acid sequences with their active sites for predicting and analyzing umami peptides. A training dataset of 204 peptides was manually collected from the literature, which randomly contained 168 of the above 188 umami peptides (in Section 3.2) and 36 non-umami peptides (Table S1 & Table S2). Based on the peptide sequence information and active site data, we study these peptides and divide their key aspects into four groups: (a) the preferred amino acid residues including D or E; (b) the preferred active site including D, E, N, Q, R, K, G, S, A, H and V; (c) the opposed residues including I, L, Y, and F and (d) the opposed active site F.

Decision trees uses the common tree visualization techniques to visualize information in the structure to display detailed information[32]. Here, we extract the key features from the dataset of 204 peptides and propose a decision tree in Figure 6 for better readability and understandability of the decision rule of umami peptide. Undoubtedly, these features vary to an extent according to the length of peptide, and the number of feature parameters gradually increases as the length of the peptide increases. Thereinto, it can be noted that the active sites in ligands are of comparable importance in this rule. As an initial test of the model based on predictive interactions. We trained the classification objective into four types, respectively. To verify the effectiveness of the proposed rule, we test on a dataset of 20 umami peptides which is chosen randomly from a set of 188 peptides above mentioned in Section 3.2. Although this rule does not identify exactly of all the peptides (92.85% peptides can be decided, and only 88.39% can be exactly decided), the expected results obtained by the unnormalized features successfully reach a rational verification accuracy (>95%) for those umami peptides and support this decision rule on umami (Fig. 7). It could be stated that this decision rule afforded high discriminating ability for distinguishing umami peptides from non-umami peptides. Thus, it can predict candidate umami peptides from various food sources.

**Fig. 6.**
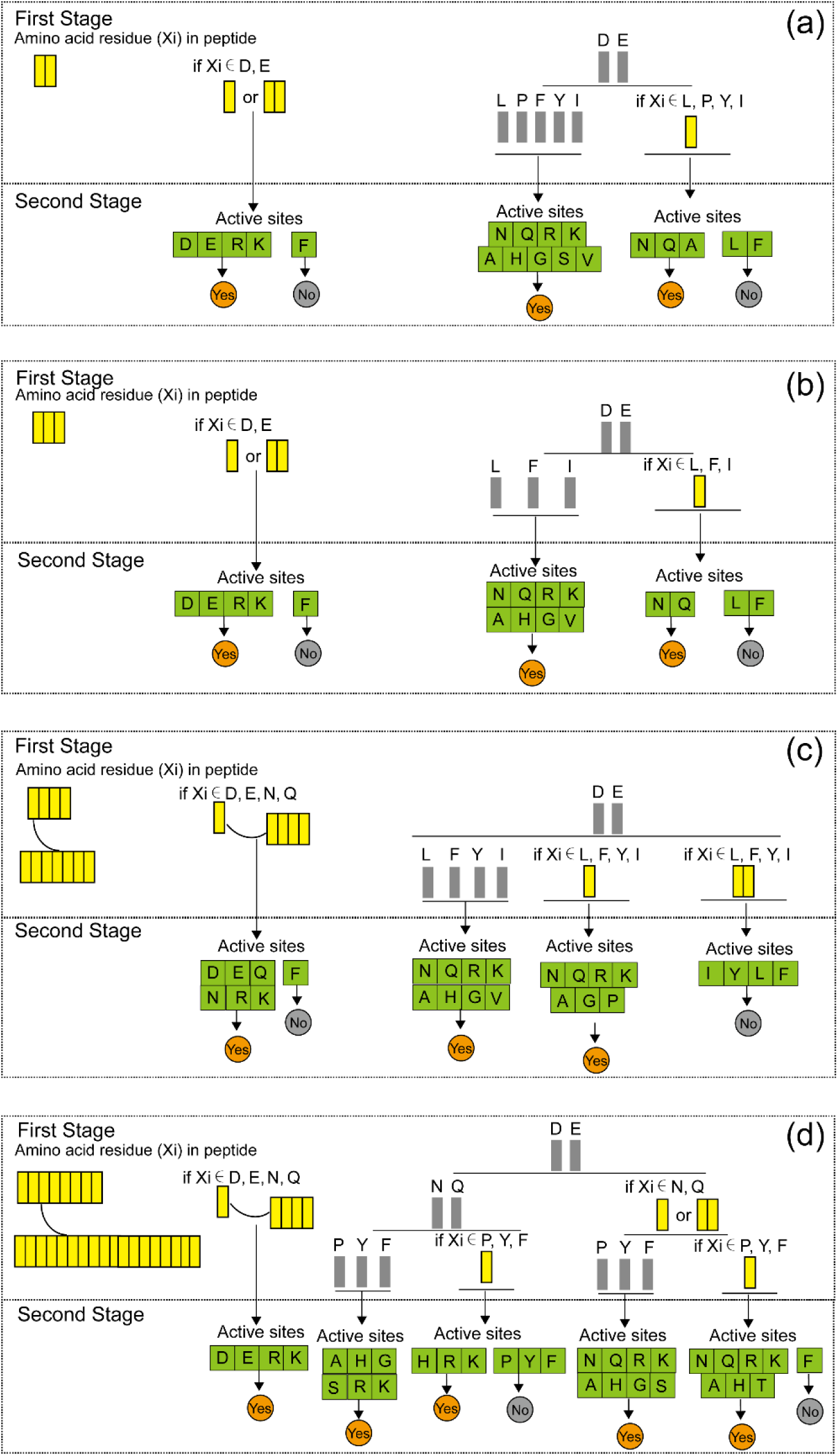
A detailed view on a decision rule in umami peptides based on their sequence information and the active sites (a): Umami peptides with 2 amino acid residues; (a): Umami peptides with 3 amino acid residues; (c): Umami peptides with 4-7 amino acid residues; (d): Umami peptides with >7 amino acid residues.

**Fig. 7.**
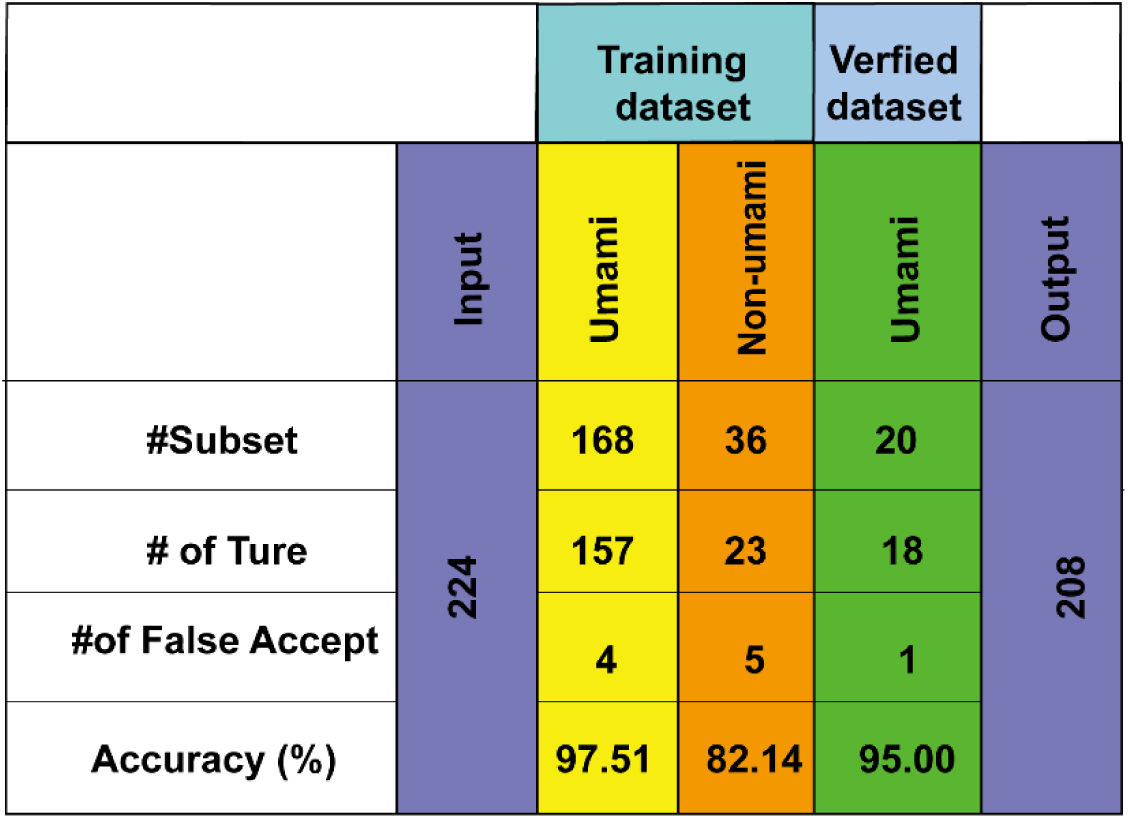
Analysis of the number of the correct results made by the proposed decision rule in umami/non-umami peptides datasets.

## Conclusion

We analyzed the sequences information and active sites of the known 205 umami peptides, and the potential binding modes of umami peptides to T1R1 were conceived by a series of the identified key binding residues in T1R1-peptide complex. Based on the sequences and active sites information of peptides, a novel decision rule for umami peptides was proposed. The results reached a rational accuracy (>95%), which further verified and supported this decision rule. It could be concluded that this decision rule was beneficial for umami peptide identification. In the meantime, the active sites of the peptide will be further utilized for interpretation of the potential physicochemical properties that are essential for the prediction of umami peptides. This is also essential for a better understanding of the umami mechanism of peptides and may open up new possibilities for the development of umami industry.

## Acknowledgments

We thank Lingkun Luo for help with data analysis and management. This work was supported by the National Natural Science Foundation of China (Grant No. 31901813, 31972198).

## Author contributions

Wenli Wang conceived the project, designed experiments, performed experiments, analysed data, wrote the paper. Zhiyong Cui performed experiments, analysed data. Menghua Ning performed experiments. Yuan Liu conceived and supervised the project, provided funding, wrote the paper.

## Appendix A. Supplementary data

**Supplementary data (Table S1)** to this article can be found online at

## References

[1] I.E. Hartley, D.G. Liem, R. Keast, Umami as an ‘Alimentary’ Taste. A New Perspective on Taste Classification, Nutrients 11(1) (2019).

[2] X.D. Li, L. Staszewski, H. Xu, K. Durick, M. Zoller, E. Adler, Human receptors for sweet and umami taste, P Natl Acad Sci USA 99(7) (2002) 4692–4696.

[3] W.L. Wang, X.R. Zhou, Y. Liu, Characterization and evaluation of umami taste: A review, Trac-Trend Anal Chem 127 (2020).

[4] J.A. Zhang, D. Sun-Waterhouse, G.W. Su, M.M. Zhao, New insight into umami receptor, umami/umami-enhancing peptides and their derivatives: A review, Trends Food Sci Tech 88 (2019) 429–438.

[5] Y. Zhang, X.C. Gao, D.D. Pan, Z.G. Zhang, T.Q. Zhou, Y.L. Dang, Isolation, characterization and molecular docking of novel umami and umami-enhancing peptides from Ruditapes philippinarum, Food Chemistry 343 (2021).

[6] J.M. Liang, L.L. Chen, Y.N. Li, X.J. Hu, Isolation and identification of umami-flavored peptides from Leccinum extremiorientale and their taste characteristic, J Food Process Pres (2021).

[7] Z. Liu, Y. Zhu, W. Wang, X. Zhou, G. Chen, Y. Liu, Seven novel umami peptides from Takifugu rubripes and their taste characteristics, Food Chemistry 330(20) (2020) 31066–9.

[8] P. Charoenkwan, J. Yana, C. Nantasenamat, M. Hasan, W. Shoombuatong, iUmami-SCM: A Novel Sequence-Based Predictor for Prediction and Analysis of Umami Peptides Using a Scoring Card Method with Propensity Scores of Dipeptides, Journal of Chemical Information and Modeling (2020).

[9] X. Yu, L. Zhang, X. Miao, Y. Li, Y. Liu, The structure features of umami hexapeptides for the T1R1/T1R3 receptor, Food Chem 221(2017) (2017) 599–605.

[10] H. Liu, L.T. Da, Y. Liu, Understanding the molecular mechanism of umami recognition by T1R1-T1R3 using molecular dynamics simulations, Biochem Bioph Res Co 514(3) (2019) 967–973.

[11] N. Nuemket, N. Yasui, Y. Kusakabe, Y. Nomura, N. Atsumi, S. Akiyama, E. Nango, Y. Kato, M.K. Kaneko, J. Takagi, M. Hosotani, A. Yamashita, Structural basis for perception of diverse chemical substances by T1r taste receptors, Nat Commun 8 (2017).

[12] J.J.L. Cascales, S.D.O. Costa, B.L. de Groot, D.E. Walters, Binding of glutamate to the umami receptor, Biophys Chem 152(1-3) (2010) 139–144.

[13] Y.L. Dang, L. Hao, J.X. Cao, Y.Y. Sun, X.Q. Zeng, Z. Wu, D.D. Pan, Molecular docking and simulation of the synergistic effect between umami peptides, monosodium glutamate and taste receptor T1R1/T1R3, Food Chem 271 (2019) 697–706.

[14] F. Zhang, B. Klebansky, R.M. Fine, H. Xu, A. Pronin, H.T. Liu, C. Tachdjian, X.D. Li, Molecular mechanism for the umami taste synergism, P Natl Acad Sci USA 105(52) (2008) 20930–20934.

[15] Y. Toda, T. Nakagita, T. Hayakawa, S. Okada, M. Narukawa, H. Imai, Y. Ishimaru, T. Misaka, Two Distinct Determinants of Ligand Specificity in T1R1/T1R3 (the Umami Taste Receptor), J Biol Chem 288(52) (2013) 36863–36877.

[16] W. Zhu, W. He, F. Wang, Y. Bu, X. Li, J. Li Prediction, molecular docking and identification of novel umami hexapeptides derived from Atlantic cod (Gadus morhua), International Journal of Food Science & Technology (2020).

[17] Z.P. Yu, L.X. Kang, W.Z. Zhao, S.J. Wu, L. Ding, F.P. Zheng, J.B. Liu, J.R. Li, Identification of novel umami peptides from myosin via homology modeling and molecular docking, Food Chemistry 344 (2021).

[18] M.N.G. Amin, J. Kusnadi, J.L. Hsu, R.J. Doerksen, T.C. Huang, Identification of a novel umami peptide in tempeh (Indonesian fermented soybean) and its binding mechanism to the umami receptor T1R, Food Chem 333 (2020) 127411.

[19] T.H. Wang, C.H. Hsiao, S.H. Chen, C.M. Hsiao, L.Y. Chen, G.M. Li, B.C. Hsueh, A DFT study on structures, frontier molecular orbitals and UV-vis spectra of [M(L)(N-3)(C7H5N)(PPh3)] (M = Ru and Fe; L = Tp and Cp), J Organomet Chem 791 (2015) 72–81.

[20] S. Damak, M.Q. Rong, K. Yasumatsu, Z. Kokrashvili, V. Varadarajan, S.Y. Zou, P.H. Jiang, Y. Ninomiya, R.F. Margolskee, Detection of sweet and umami taste in the absence of taste receptor T1r3, Science 301(5634) (2003) 850–853.

[21] H.N. Lioe, K. Takara, M. Yasuda, Evaluation of peptide contribution to the intense umami taste of Japanese soy sauces, Journal of Food Science 71(3) (2006) S277–S283.

[22] J.A. Zhang, M.M. Zhao, G.W. Su, L.Z. Lin, Identification and taste characteristics of novel umami and umami-enhancing peptides separated from peanut protein isolate hydrolysate by consecutive chromatography and UPLC-ESI-QTOF-MS/MS, Food Chemistry 278 (2019) 674–682.

[23] B.C. Popere, A.M. Della Pelle, A. Poe, G. Balaji, S. Thayumanavan, Predictably tuning the frontier molecular orbital energy levels of panchromatic low band gap BODIPY-based conjugated polymers, Chem Sci 3(10) (2012) 3093–3102.

[24] N.L. Zhang, W.L. Wang, B. Li, Y. Liu, Non-volatile taste active compounds and umami evaluation in two aquacultured pufferfish (Takifugu obscurus and Takifugu rubripes), Food Biosci 32 (2019).

[25] N. Ishibashi, I. Ono, K. Kato, T. Shigenaga, I. Shinoda, H. Okai, S. Fukui, Role of the Hydrophobic Amino Acid Residue in the Bitterness of Peptides, Agricultural & Biological Chemistry 52(1) (1988) 91–94.

[26] P. Charoenkwan, J. Yana, N. Schaduangrat, C. Nantasenamat, M.M. Hasan, W. Shoombuatong, iBitter-SCM: Identi fication and characterization of bitter peptides using a scoring card method with propensity scores of dipeptides, Genomics 112(4) (2020) 2813–2822.

[27] Y.L. Huang, D.Q. Lu, H. Liu, S.Y. Liu, S. Jiang, G.C. Pang, Y. Liu, Preliminary research on the receptor-ligand recognition mechanism of umami by an hT1R1 biosensor, Food Funct 10(3) (2019) 1280–1287.

[28] P. Charoenkwan, J. Yana, C. Nantasenamat, M. Hasan, W. Shoombuatong, iUmami-SCM: A Novel Sequence-Based Predictor for Prediction and Analysis of Umami Peptides Using a Scoring Card Method with Propensity Scores of Dipeptides, Journal of Chemical Information and Modeling 60(12) (2020) 6666–6678.

[29] A. Dunkel, T. Hofmann, A. Di Pizio, In Silico Investigation of Bitter Hop-Derived Compounds and Their Cognate Bitter Taste Receptors, J Agr Food Chem 68(38) (2020) 10414–10423.

[30] A.D.-W. Eitan Margulis, Robert S. Ives, Sara Jaffari, Karsten Siems, Masha Y. Niv, Intense bitterness of molecules: Machine learning for expediting drug discovery, Computational and Structural Biotechnology Journal 19 (2021) 568–576.

[31] A. Goel, K. Gajula, R. Gupta, B. Rai, In-silico screening of database for finding potential sweet molecules: A combined data and structure based modeling approach, Food Chemistry 343 (2021).

[32] T.D. Nguyen, T.B. Ho, H. Shimodaira, Interactive visualization in mining large decision trees, Lect Notes Artif Int 1805 (2000) 345–348.

